# Proteome remodeling in the *Mycobacterium tuberculosis* PknG knockout: molecular evidence for the role of this kinase in cell envelope biogenesis and hypoxia response

**DOI:** 10.1101/2021.03.16.435434

**Authors:** Analía Lima, Alejandro Leyva, Bernardina Rivera, María Magdalena Portela, Magdalena Gil, Alessandro Cascioferro, María-Natalia Lisa, Annemarie Wehenkel, Marco Bellinzoni, Paulo C. Carvalho, Carlos Batthyány, María N. Alvarez, Roland Brosch, Pedro M. Alzari, Rosario Durán

## Abstract

*Mycobacterium tuberculosis*, the etiological agent of tuberculosis, is among the deadliest human pathogens. One of *M. tuberculosis*’s pathogenic hallmarks is its ability to persist in a dormant state in the host for long periods, reinitiating the infectious cycle when favorable environmental conditions are found. Thus, it is not surprising that this pathogen has developed different mechanisms to withstand the stressful conditions found in the host. In particular, the Ser/Thr protein kinase PknG has gained special relevance since it regulates nitrogen metabolism and facilitates bacterial survival inside macrophages. Nevertheless, the molecular mechanisms underlying these effects are far from being elucidated. To further investigate these issues, we performed quantitative proteomics analyses of protein extracts from *M. tuberculosis* H37Rv and a mutant derivative lacking *pknG*. Our results showed that in the absence of PknG the mycobacterial proteome was remodeled since 5.7% of the proteins encoded by *M. tuberculosis* presented significant changes in its relative abundance when compared to the wild-type strain. The main biological processes affected by *pknG* deletion were the biosynthesis of cell envelope components and the response to hypoxic conditions. As many as 13 DosR-regulated proteins were underrepresented in the *pknG* deletion mutant, including the distinctive Hrp-1, which was found to be 12-fold decreased according to Parallel Reaction Monitoring experiments. Altogether, the results presented here allow us to postulate that PknG regulation of bacterial adaptation to stress conditions might be an important mechanism underlying its reported effect on intracellular bacterial survival.

## INTRODUCTION

Tuberculosis (TB) is a pulmonary disease caused by *Mycobacterium tuberculosis* that remains a major global health problem, being responsible for around 1.4 M deaths worldwide during 2019 (1). Clinically, it can be an active (transmissible and symptomatic), sub-clinical (transmissible but without symptoms), or latent disease (non-transmissible and asymptomatic) (2). It is estimated that around a quarter of the human population is affected by the latent form of TB, which constitutes a large reservoir for the pathogen (1). The main pathogenic characteristic of *M. tuberculosis* is its ability to arrest phagosome maturation, allowing bacterial survival and replication inside relatively harmless vacuole (3). Macrophage infection triggers a localized pro-inflammatory response that leads to the formation of granulomas, a hallmark of TB, where *M. tuberculosis* can persist for decades in a protected environment, mostly in a dormant state under hypoxic conditions (4). In the human host, *M. tuberculosis* encounters many different ecological niches and exhibits various physiological states. Thus, it is not surprising that *M. tuberculosis* has developed some very peculiar structural traits and a diversity of regulatory and metabolic capabilities to cope with these different stress conditions.

The impermeable cell envelope is a distinctive characteristic of mycobacteria. It is composed of a central core of peptidoglycan covalently attached to arabinogalactans esterified with mycolic acids, which constitute the inner leaflet of the mycomembrane. Externally, there is an additional layer composed of non-covalently attached (glyco)lipids that are very important for pathogenesis and the host-bacterium interaction (5, 6). Additionally, *M. tuberculosis* has developed a variety of strategies to withstand host’s stress conditions: an unusually high number of toxin-antitoxin systems (TA systems) (7), different mechanisms to resist the host oxidative attack (8) and to adapt to low nutrient concentration, as well as low oxygen tension (9, 10) and protein phosphorylation systems based on bacterial “eukaryotic-like” Ser/Thr protein kinases (STPKs) and two-component systems (TCS). Concerning the latter, the DosSR two-component system is crucial for the bacterial adaptation to the redox status and to low oxygen concentrations through the induction of around fifty genes comprised within the mycobacterial Dormancy Survival Regulon (DosR regulon, also known as DevR regulon) (11–14).

Among STPKs, PknG was shown to play an important role in bacterial survival within the host, in bacterial metabolism, and in pathogenesis (15–19). PknG participates in phagosome maturation inhibition promoting *M. tuberculosis* survival inside macrophages (15) and facilitates bacterial growth under *in vitro* stress conditions, such as nutrient deprivation, acid stress, and hypoxia (20, 21). PknG additionally contributes to the intrinsic antibiotic resistance of pathogenic mycobacteria, as the deletion of *pknG* caused a multidrug sensitive phenotype (22). Moreover, the activity of several enzymes that participate in glutamate metabolism is regulated by PknG through the phosphorylation of the FHA-domain-containing substrate GarA (17). PknG also phosphorylates the ribosomal protein L13, triggering the regulation of the activity of the Nudix hydrolase RenU (23). Phosphoproteomics, interactomics and protein array studies have allowed to expand the list of putative mycobacterial PknG substrates (24–26), which currently comprises up to 31 proteins phosphorylated by this kinase either *in vivo* or *in vitro* (27). However, the physiological relevance of these phosphorylation events and their possible relationship to the proposed roles of the kinase in bacterial survival within the host are not yet fully understood.

The objective of this work was to contribute to the elucidation of the biological processes regulated by PknG in mycobacteria. For this purpose, we employed two complementary quantitative proteomic approaches, 2D-DIGE (Differential In Gel Electrophoresis) and label-free LC-MS/MS, to analyze protein extracts from *M. tuberculosis* H37Rv and a mutant strain lacking PknG. Both approaches jointly shortlisted around 300 differentially abundant proteins. We show that *M. tuberculosis* H37Rv Δ*pknG* presents altered levels of proteins involved in response to hypoxia, TA systems and synthesis of the core as well as outer layer lipids of the cell wall. Altogether, these results allow us to postulate that the regulation of the expression levels of proteins that are essential for bacterial fitness to host stress conditions is an important mechanism supporting the reported effects of PknG on bacterial survival within the host.

## MATERIAL AND METHODS

### Mycobacterial cultures and protein extract preparation

The *Mycobacterium tuberculosi*s *pknG* null mutant strain (Δ*pknG*) was kindly provided by Josef Av-Gay (18). Wild-type *M. tuberculosis* H37Rv (WT) and Δ*pknG* strains were cultured in 50 mL of Middlebrook 7H9 medium supplemented with 0.05% Tween® 80, albumin-dextrose and asparagine (BD Biosciences) until early-logarithmic phase as previously described (19). Mycobacterial cells were washed with PBS buffer, resuspended in minimum medium supplemented with 10 mM asparagine and cultured for five additional days. Cells were harvested by centrifugation (3000 g for 10 min at 4 °C), washed in PBS buffer containing the Complete EDTA-free Protease Inhibitor Cocktail (Roche) in the amount recommended by the supplier. To prepare whole cell lysates, an equal amount of acid-washed glass beads (≤ 106 µm, Sigma) was added to the cell suspension and the system was vortexed at maximum speed for 10 min. Cell debris and beads were removed by centrifugation at 1,000 g for 5 min at 4 °C and lysates filtered through 0.22 μm PVDF membranes and stored at −80 °C for further analysis. Protein quantification was performed by gel densitometry measurements using the 1D analysis module of the ImageQuant TL software (v8.1) and the LMW-SDS Marker Kit (GE Healthcare) as standard. Protein extracts for the WT and Δ*pknG M. tuberculosis* strains were prepared in triplicate.

### Differential In-Gel Electrophoresis (DIGE)

A former DIGE analysis comparing *M. tuberculosis* H37Rv and Δ*pknG* was focused on two proteins that presented pI variations compatible with protein phosphorylation (24). In the present work DIGE analysis on the same gels set was performed with a global proteome perspective.

Differential and quantitative analyses between *M. tuberculosis* strains WT and Δ*pknG* were carried out for three biological replicates employing the Ettan DIGE System (GE Healthcare) following manufacturer’s recommendations (28) and as described (24). Briefly, 100 μg of each protein extract were concentrated using the 2D Clean-up kit (GE Healthcare) and then resuspended in 30 mM Tris pH 8.5, 7 M urea, 2 M thiourea, 4% CHAPS. Twenty-five μg of each WT and Δ*pknG* samples were combined to prepare the internal standard (STD) used for 2D-DIGE image matching, spot volume normalization and abundance change determinations.

Fifty μg of each STD, WT, and Δ*pknG* samples were differentially labeled with cyanine dyes Cy2, Cy3, and Cy5 following the manufacturer’s instructions (GE Healthcare) (28). To compensate for any labeling bias, Cy3 and Cy5 were alternatively employed to label the WT and Δ*pknG* proteomes, while Cy2 was exclusively used for labeling the STD. Differentially labeled WT, Δ*pknG*, and STD samples were mixed and the rehydration solution (7 M urea, 2 M thiourea, 4% CHAPS, 0.5% IPG Buffer 4-7 [GE Healthcare]) was added before overnight IPG-strips passive rehydration (pH gradient 4-7, 13 cm).

Isoelectric focusing was performed in an IPGphor Unit (Pharmacia Biotech) employing the previously recommended voltage profile (29). Then, IPG-strips were treated for 15 min in equilibration buffer (6 M urea, 75 mM Tris–HCl pH 8.8, 29.3% glycerol, 2% SDS, 0.002% bromophenol blue) with the addition of 10 mg/mL DTT and next in the same buffer with 25 mg/mL iodoacetamide. The second-dimension separation was carried on a 12.5% SDS-PAGE in a SE 600 Ruby Standard Dual Cooled Vertical Unit (GE Healthcare) at 20 °C. Images from gels were obtained employing a Typhoon FLA 9500 variable mode laser scanner (GE Healthcare) at a resolution of 100 μm using the laser wavelength and the filter settings indicated for each dye (30). The photomultiplier tube voltage was adjusted on each channel to give maximum pixel values below saturation levels (60,000-90,000 counts).

The analysis of images was performed using DeCyder^TM^ 2D software (v7.2) (GE Healthcare). The DeCyder’s Differential In-gel Analysis module (DIA) was employed for spots co-detection, quantification by normalization, and ratio calculation. The DeCyder’s Biological Variation Analysis module (BVA) was utilized for gel-to-gel spot matching and statistical analysis, allowing quantitative comparisons of spot volumes across multiple gels. An unpaired Student’s t-test was used to assess significant changes. Spots displaying significant differences (*p-*value ≤ 0.05) and a fold-change greater than 25% were selected for further analyses.

### Protein identification by MALDI-TOF/TOF MS

Differential spots were matched to the silver-stained master gel and were picked and processed for MALDI-TOF/TOF analysis following previously reported protocols (31, 32).

Mass spectra of peptides mixtures were acquired in a 4800 MALDI TOF/TOF instrument (ABiSciex, USA) in positive ion reflector mode. Mass spectra were calibrated using a mixture of peptides standard (Applied Biosystems) and trypsin autolysis products. Some peptides from all protein spots were selected for MS/MS analyses using the following settings: 8 kV and 15 kV for sources 1 and 2, respectively.

Protein identification was performed by database searching of acquired *m/z* values employing the MASCOT search engine (Matrix Science http://www.matrixscience.com/search_form_select.html) in the Sequence Query mode, using a database from NCBI (20170811), and applying the following search parameters: monoisotopic mass tolerance, 0.05 Da; fragment mass tolerance, 0.5 Da; Met oxidation and Ser/Thr/Tyr phosphorylation as variable modifications, Cys carbamidomethylation as fixed modification, and allowance of one missed tryptic cleavage. Protein mass was unrestricted and taxonomy was limited to *Mycobacterium tuberculosis* complex. Significant protein scores (*p*-value < 0.05) were used as criteria for confident identification.

### nanoLC-MS/MS analysis and protein identification

Total protein extracts (20 μg) from WT and Δ*pknG* samples (in triplicates) were reduced with 10 mM DTT at 56 °C for 60 min and then alkylated with 55 mM iodoacetamide at room temperature for 45 min. The samples were separated by SDS-PAGE using precast 4%-12% gradient gels (NuPAGE, MES System, Invitrogen) and stained with Coomassie Brilliant Blue G-250. Each lane was excised into ten slices that were destained by incubation with 0.2 M ammonium bicarbonate/ACN (1:1) for 1 h at room temperature with agitation. *In-gel* proteolytic digestion and peptide extraction were performed as described earlier (24). Peptide samples were vacuum dried, resuspended in 0.1% formic acid and injected into a nano-HPLC system (EASY-nLC 1000, Thermo Scientific) equipped with a reverse-phase column (EASY-Spray column, 50 cm × 75 µm ID, PepMap RSLC C18, 2 µm, Thermo Scientific). Peptides separation was performed at a constant flow rate of 250 nL/min and using a gradient from 0% to 50% of mobile phase B (mobile phase A: 0.1% formic acid, mobile phase B: 0.1% formic acid in acetonitrile) over 100 min. Peptide analysis was performed in an LTQ Velos nano-ESI linear ion trap equipment (Thermo Scientific) in a data-dependent acquisition mode. Xcalibur 2.1 was used for data acquisition in two steps: 1. acquisition of full MS scan in the positive ion mode with *m/z* between 300 and 1800 Da, 2. sequential fragmentation of the ten most intense ions with a normalized collision energy of 35, an isolation width of 2 *m/z*. The activation Q was set on 0.25, the activation time on 15 ms, and a dynamic exclusion time of 30 s. MS source parameters were set as follows: 2.3 kV electrospray voltage and 260 °C capillary temperature.

PatternLab for Proteomics (version 4.0.0.74) (33) was employed to generate a target-decoy database using sequences from *M. tuberculosis* H37Rv (taxon identifier: 83332; 3993 sequences) downloaded from the UniProt consortium in October, 2017. In addition, 127 common mass spectrometry contaminants were included (33) giving rise to a target-reverse database with 8228 entries.

The Comet search engine was operated using the following parameters: trypsin as proteolytic enzyme with full specificity; oxidation of Met and phosphorylation on Ser/Thr/Tyr as variable modifications, carbamidomethylation of Cys as fixed modification; and 800 ppm of tolerance from the measured precursor *m/z*. XCorr and Z-Score were used as the primary and secondary search engine scores, respectively. Raw data is available in the ProteomeXchange Consortium via the PRIDE partner repository (34) with the identifier PXD023975.

Peptide spectrum matches were filtered using the Search Engine Processor (SEPro) and acceptable false discovery rate (FDR) criteria were set on ≤ 1% at the protein level, and ≤ 2% at the peptide level. The actual FDR for each file searched is depicted in Supplementary Table 1. PatternLab’s statistical model for the Approximately Area Proportional Venn Diagram module was used to compare conditions and determine proteins that are likely to be exclusively detected in each situation due to differences in its abundance (*p*-value < 0.05). The Bayesian model integrated into PatternLab for Proteomics (35) considers quantitative data and the number of appearances in different biological replicates to assign *p*-values. PatternLab’s T-Fold module was used to detect proteins present in both conditions at significantly different levels by spectral count analysis. Buzios and Clustergram modules were used to perform a Principal Component Analysis and a Heatmap, respectively (33).

### Parallel Reaction Monitoring (PRM) Targeted MS

Some proteins, found to display differential abundances by discovery proteomics, belonging to the metabolism of the outermost lipids of the mycobacterial cell envelope (Mas and Msl3) or involved in the response to hypoxia (Icl, Ald, Lat, Rv2030c, Acg, DevR, hspX, Hrp-1) were chosen to be validated by tier 3 targeted proteomic analysis.

For parallel reaction monitoring (PRM) analysis, 20 μg of WT and Δ*pknG* samples were run 1 cm on a resolving SDS-PAGE and processed for mass spectrometry analysis as described above, using 20 μL of mobile phase A to resuspend them.

To generate the spectral library, 5 μL of each sample were mixed and separated using a nano-HPLC (UltiMate 3000, Thermo) coupled to a Q-Orbitrap mass spectrometer (Q-Exactive Plus, Thermo). Tryptic peptides (5 μg) were injected into an Acclaim PepMap^TM^ 100 C18 nano-trap column (75 μm x 2 cm, 3 μm particle size, Thermo) and separated in a 75 μm x 50 cm, PepMap^TM^ RSLC C18 analytical column (2 μm particle size, 100 Å, Thermo) at a constant flow rate of 200 nL/min and 40 °C. The column was equilibrated with 1% of mobile phase B, and the elution was performed using a gradient from 1% to 50% of mobile phase B over 180 min and 50% to 99% over 15 min. The ion spray voltage setting was 1.7 kV, the capillary temperature was set at 250 °C and S-lens RF level at 50. Mass analysis was performed in a data-dependent mode in two-step: 1. acquisition of full MS scan in a 200 to 2000 *m/z* range; 2. fragmentation of the 12 most intense ions in each segment by HCD using a stepped normalized collision energy of 25, 30, and 35. The following settings were used for full MS scans: a resolution of 70,000 at 200 *m/z*, an AGC target value of 1E06, and a maximum ion injection time of 100 ms. For MS/MS acquisition the resolution was 17,500 at 200 *m/z*, AGC target value of 1E05, and maximum injection time of 50 ms. Precursor ions with unassigned, single, and higher than five charges were excluded. The dynamic exclusion time was set at 10 s. Two technical replicates of the sample mix were used to generate the library. Thermo raw files were searched against the target-decoy *M. tuberculosis* H37Rv database as described above but using PatternLab (v5). Search was performed with 40 ppm for precursor mass accuracy and results were afterwards filtered using 5 ppm error tolerance. The result file was saved in SSL format as .raw file (Thermo), with carbamidomethylation of Cys as a fixed modification, to interface with Skyline software.

Taking into account the *M. tuberculosis* H37Rv background proteome downloaded from the UniProt server (www.uniprot.org) and the generated spectral library data, 23 peptides derived from the 11 chosen proteins were selected using Skyline (v. 20.1.0.155). An unscheduled isolation list was generated, loaded into Thermo Xcalibur 4.0.27.19 and used to carry out a PRM analysis on the Q-Exactive Plus mass spectrometer. Five μg of WT and Δ*pknG* samples were injected in duplicates and peptides were separated using the same gradient used for spectral library generation. The mass spectrometer settings were as follows: positive polarity, a resolution of 17,500 at 200 *m/z*, AGC target value of 2E04, maximum injection time of 50 ms, 2.0 *m/z* of isolation window, stepped normalized collision energy of 25, 30, and 35. The list of the targeted peptides in the PRM analysis is shown in Supplementary Table 2. The mass spectrometry proteomics data have been deposited to the ProteomeXchange Consortium via th PRIDE partner repository (34) with the dataset identifier PXD023956.

Resulting PRM raw data was extracted and imported into Skyline software (v20.2) for analysis. Data were manually refined using the spectral library as a reference, taking into account the dotp values for each peptide, and the comparison of peak areas for each transition. For quantitative comparative analyses between WT and Δ*pknG* peptides, the sum of the peak areas of the transitions of each peptide and the equalization to medians were employed. Peptides with a significant difference showing an adjusted *p*-value < 0.05 were considered.

### Experimental Design and Statistical Rationale

Protein extracts from *M. tuberculosis* H37Rv and from the same strain in which the *pknG* gene was inactivated by allelic exchange were used to study the effect of this kinase deletion on global proteome profile. Three biological replicates per condition were used for discovery proteomics experiments, while two biological replicates were examined for PRM experiments. In this last case technical replicates of each sample were analyzed.

### Bioinformatics Analyses

The functional classification of identified proteins was performed using the information provided by the Mycobrowser server (https://mycobrowser.epfl.ch/).

A statistical overrepresentation test was performed using the Panther Server Classification System (http://pantherdb.org) version 15.0, released 2020-02-14 (36). The annotation data set “GO biological processes complete” (released 2020-07-28) was used. The release date of the GO ontology dataset was 2020-10-09. Protein showing significant abundance changes, by both 2D-DIGE and label-free LC-MS/MS approaches, were used for the analysis. Proteins that only show changes in proteoforms patterns were excluded. The *M.tuberculosis* database was used as a reference List.

## RESULTS

To identify proteins and processes altered in a *M. tuberculosis pknG* deletion mutant that could be responsible for the previously observed phenotypes in bacterial survival, we compared the proteomes of wild-type *M. tuberculosis* H37Rv (WT) and Δ*pknG* using two complementary quantitative approaches: 2D-DIGE and label free LC-MS/MS analysis.

### 2D-DIGE analysis revealed that proteins involved in lipid metabolism and stress adaptation are altered in M. tuberculosis ΔpknG

Protein extracts from *M. tuberculosis* WT and Δ*pknG* cells were prepared in triplicates and analyzed by 2D-DIGE (24). A representative 2D-DIGE gel image, showing WT proteins labeled with Cy5 dye (red spots) and Δ*pknG* proteins labeled with Cy3 (green spots), is depicted in Figure 1.

**Figure 1:**
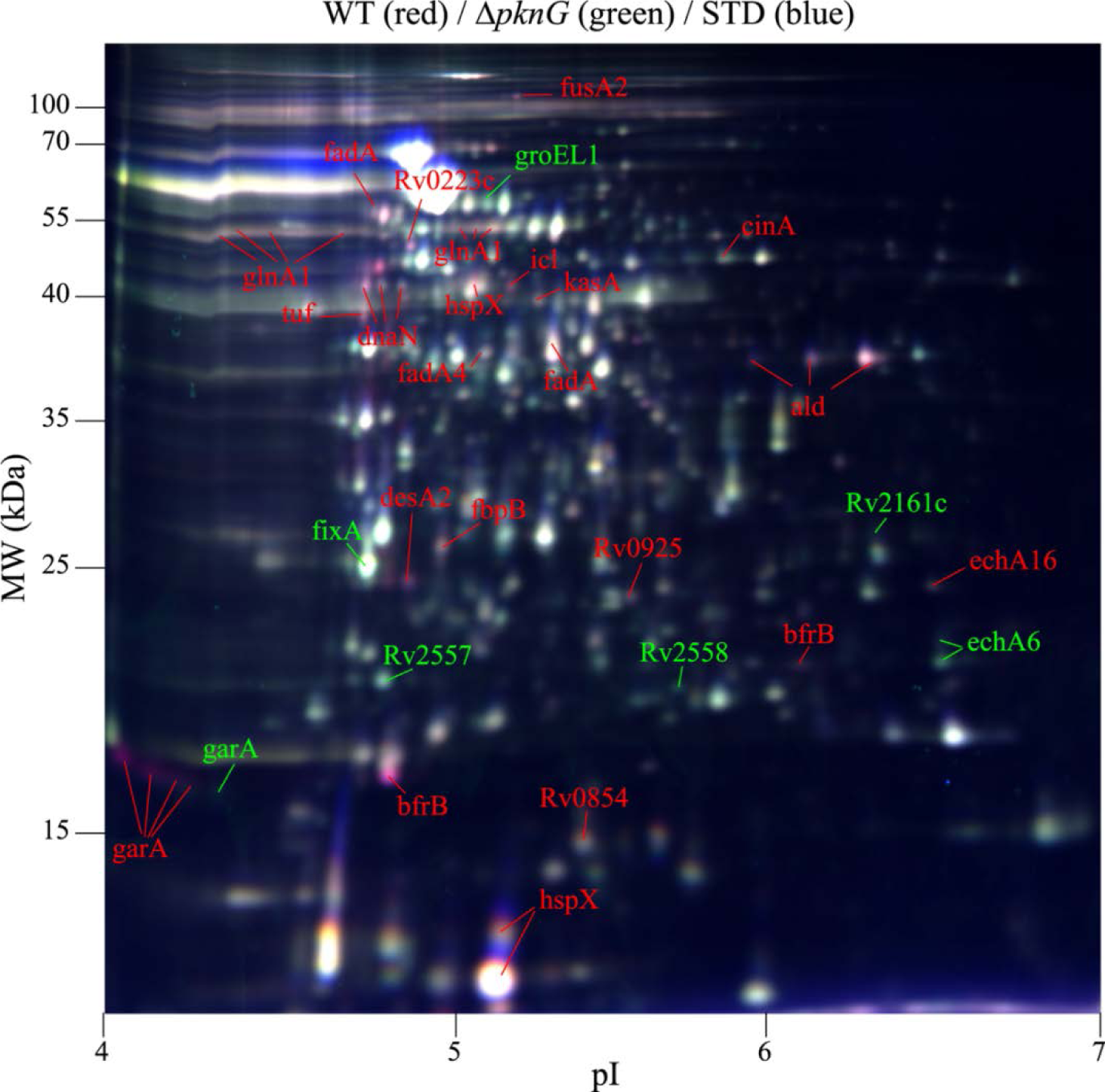
Representative 2D-DIGE image of total protein extracts from *M. tuberculosis* H37Rv WT and Δ*pknG* strains. Overlay image of WT (labeled with Cy5, red), Δ*pknG* (labeled with Cy3, green), and the internal standard (STD, labeled with Cy2, blue). Identified differential spots (considering all replicates, *p*-value ≤ 0.05 and fold-changes ≥ 25%) are shown. Red and green labels indicate spots overrepresented in WT and Δ*pknG*, respectively. GarA and GlnA1 spots were previously identified (24). Details regarding 2D-DIGE analyses and protein identification values for all the other differential spot are depicted in Supplementary Figure 1 and Supplementary Table 3.

Image analysis allowed detecting 111 spots with statistically significant abundance differences between the two strains (fold-changes > 25%; *p*-value ≤ 0.05), 30 of which were overrepresented while 81 were underrepresented in Δ*pknG*. Sixty-six spots could be confidently excised from the post-stained master gel and MALDI TOT/TOF MS analysis led to the identification of 27 different proteins from 36 spots (Supplementary Figure 1). More than one protein was identified in two spots (spots 27 and 36), and therefore they were not considered for further analyses. Several differential spots were previously assigned to proteoforms of GarA and GlnA1 (24), two earlier reported substrates of PknG involved in nitrogen metabolism (17, 19, 24). Thirty-six new differential spots, indicated in Figure 1, emerged from this analysis (further details are given in Supplementary Table 3). Many of the underrepresented proteins in Δ*pknG* are involved in lipid metabolism processes, including fatty acid degradation (FadA, EchA16 and FadA4), biosynthesis of unsaturated fatty acids (DesA2), biosynthesis of mycolic acids (KasA) and triacylglycerol metabolism (FbpB). Notably, 10 underrepresented spots in the Δ*pknG* strain corresponded to proteins upregulated in hypoxic and other growth-limiting conditions that usually lead to non-replicative mycobacteria (4, 37–41). These spots corresponded to the bacterioferritin BrfB, the heat shock protein HspX, isocitrate lyase Icl, the elongation factor Tu and the secreted L-alanine dehydrogenase Ald (Figure 1 and Supplementary Table 3). Conversely, the uncharacterized proteins Rv2557 and Rv2558, which were previously reported to be overexpressed in *M. tuberculosis* under starvation conditions (42), were found to be overrepresented in the Δ*pknG* strain. Some of the identified proteins led to more than one spot, and some of them had not the expected pI and/or MW, possibly reflecting the presence of post-translational modifications.

Overall, our 2D-DIGE analysis suggested that *M. tuberculosis* Δ*pknG* presents alterations in glutamate as well as lipid metabolism and has modified levels of proteins (and/or its proteoforms) required for the bacterial adaptation to stress conditions known to induce a persistent state.

### Label-free LC-MS/MS analysis indicated that the cell envelope lipid biosynthesis, hypoxia, and other stress-related processes are altered in M. tuberculosis ΔpknG

LC-MS/MS analysis allowed identifying an average of 1513 proteins among replicates, which correspond to approximately 38% of the total proteins encoded by *M. tuberculosis* H37Rv, denoting a substantial coverage of the proteome. Supplementary Table 1 shows all proteins detected in each replicate, its UniProt accession number, number of unique peptides, sequence counts, spectrum counts, normalized spectral abundance factor (NSAF), protein sequence coverage as well as protein score. Information concerning peptides assigned to each protein (including precursor charge at maximum primary score and observed peptide modifications) is provided in Supplementary Table 4.

Using the Venn diagram’s statistic module from the PatternLab for Proteomics software (33), 59 proteins were exclusively detected in *M. tuberculosis* WT whereas 32 proteins were solely detected in Δ*pknG* (*p-*value < 0.05) (Figure 2, Supplementary Table 5). Then, we used the statistics PatternLab for Proteomics TFold module to compare the proteins present in both proteomes but with differential abundance (33). Considering proteins present in at least 4 replicates in all classes, 137 proteins were found with statistically different levels in Δ*pknG* samples when compared to WT (*q*-value ˂ 0.05) (Figure 3 and Supplementary Table 5), 45 of them were overrepresented while 92 were underrepresented in *M tuberculosis ΔpknG*.

**Figure 2:**
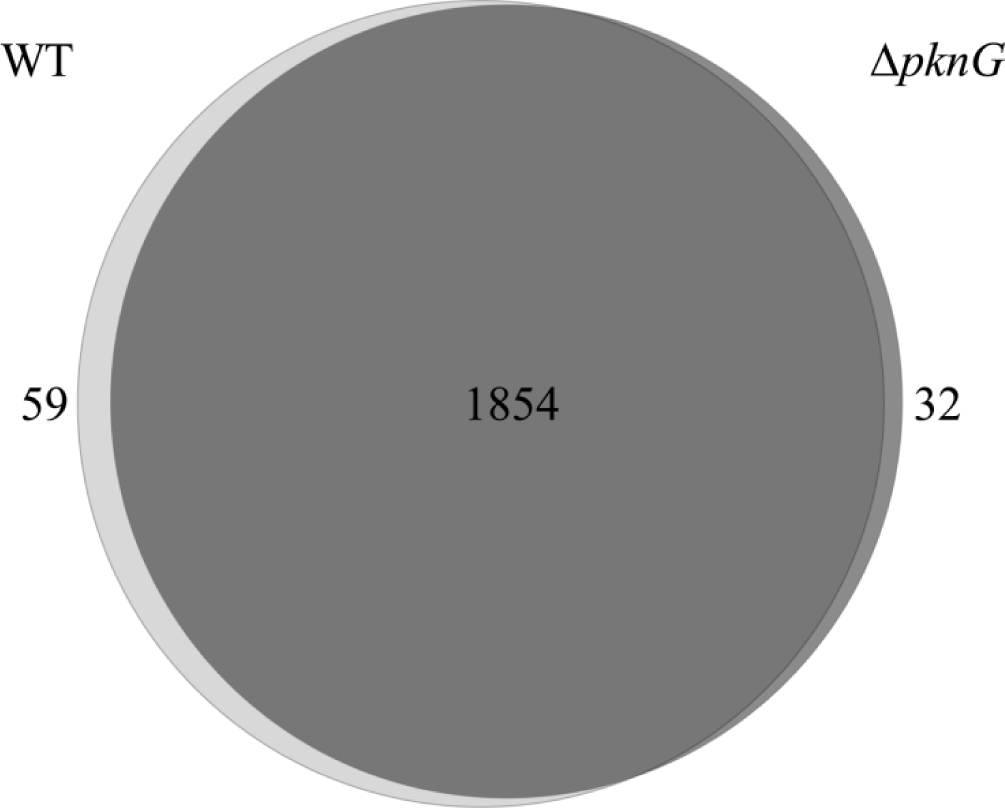
Proteins uniquely detected in *M. tuberculosis* H37Rv WT and Δ*pknG*. Venn diagram showing the number of proteins determined as statistically exclusively detected (*p*-value ˂ 0.05) in WT (59 proteins, light gray) and Δ*pknG* (32 proteins, dark gray) protein extracts, determined using the Venn diagram module from PatternLab for Proteomics software. The number of proteins identified in both strains, in at least two replicates of each, is depicted (1854 proteins). The list of proteins is detailed in Supplementary Table 5.

**Figure 3:**
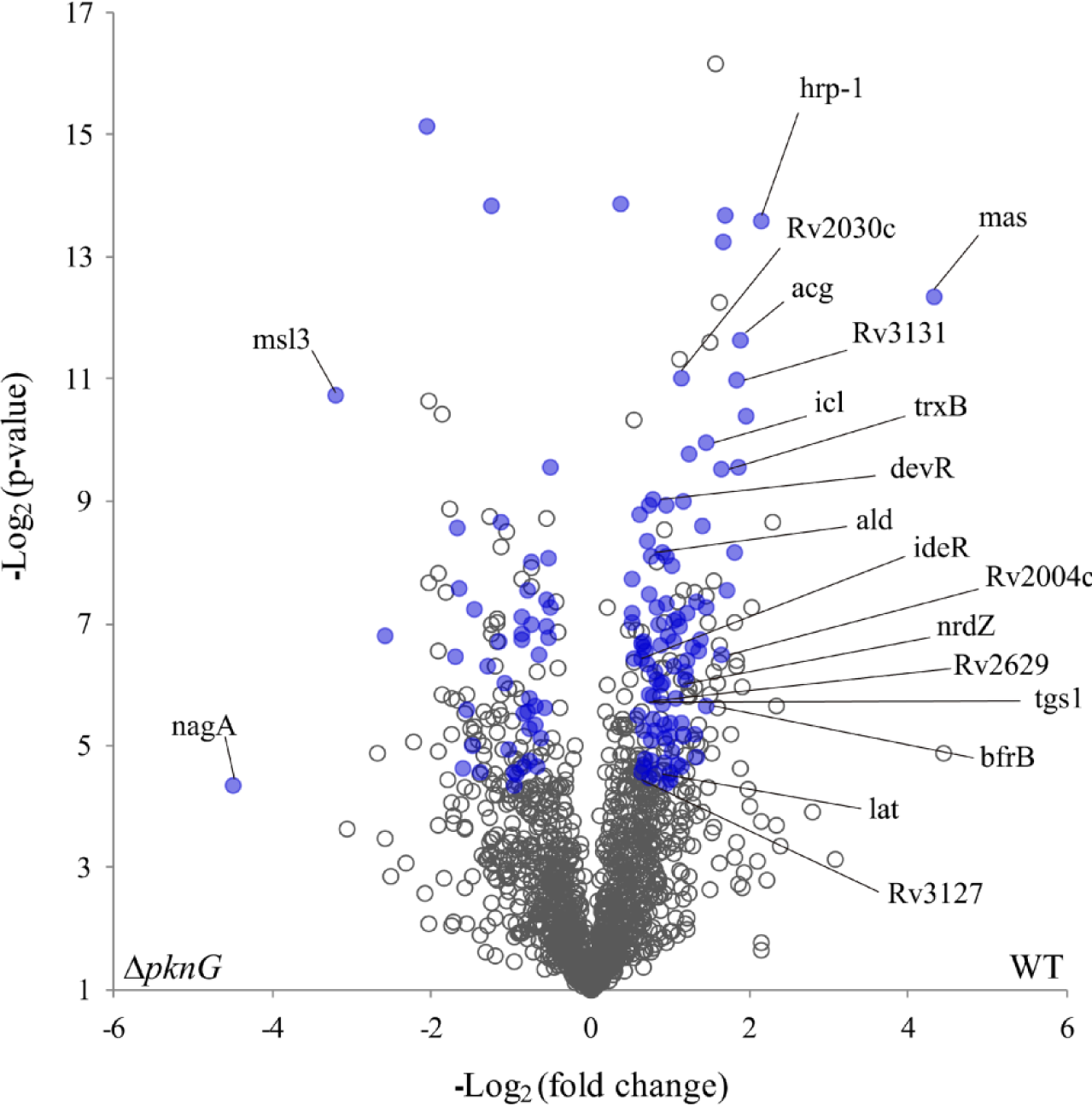
Differentially abundant proteins between *M. tuberculosis* H37Rv WT and Δ*pknG*. PatternLab’s TFold module was used to pinpoint proteins found in both conditions but showing a statistically differential abundance (BH q-value < 0.05). The volcano plot shows the Log_2_ (*p*-value) on the y-axis and the Log_2_ (fold-change) on the x-axis. Each dot in the plot represents a protein detected in at least 4 replicates of all conditions. Blue dots correspond to proteins satisfying all statistical filters and are considered differentially abundant proteins between strains. Selected differential proteins discussed in the text are labeled and all the information regarding differential proteins is depicted in Supplementary Table 5.

A principal component analysis (PCA) performed on the samples using the Búzios module of Patternlab for Proteomics showed the correct grouping of the WT and Δ*pknG* sample sets, confirming the consistency in the global observed proteomics changes (Supplementary Figure 2). In addition, hierarchical clustering classified the three replicates of the WT strain in the same hierarchical cluster but different form the one of the Δ*pknG* replicates (Supplementary Figure 3).

Not surprisingly, among the proteins exclusively detected in WT, PknG was the one identified with the highest number of spectra, followed by PhoH2, a protein comprising a TA system with RNAse activity (43). Additional TA system members were also detected exclusively in WT proteomes (the toxins VapC38, VapC37, HigB2; and the antitoxins VapB24 and MazE3).

Notably, many proteins that integrate the DosR regulon showed decreased abundance in Δ*pknG*. Three of them (the 6-phosphofructokinase PfkB, the conserved hypothetical proteins Rv2003c, and Rv3134c) were uniquely detected in the WT dataset and many others were underrepresented in Δ*pknG,* including vitamin B12-dependent ribonucleoside-diphosphate reductase NrdZ, Rv2004c, Rv2030c, the putative NAD(P)H nitroreductase Acg, the hypoxic response protein Hrp-1, Rv2629, Rv3127, the probable diacylglycerol O-acyltransferase Tgs, Rv3131 and the response regulator DosR itself (Figure 3, Table 1, Supplementary Table 5) (12, 14, 44). In a similar vein, other proteins important for the hypoxia-induced persistent state (40, 41, 45, 46) were diminished in Δ*pknG*, such as the isocitrate lyase Icl, the secreted L-alanine dehydrogenase Ald, and the probable L-lysine-epsilon aminotransferase Lat (Figure 3, Supplementary Table 5). In addition, a set of proteins involved in redox sensing and reaction to host oxidative attack also showed decreased abundance in Δ*pknG*, including IdeR (an iron-dependent response regulator that also senses redox status) (47); the bacterioferritin BfrB (48), the thioredoxin reductase TrxB and DosR, which responds to changes in both oxygen tension and redox status (10).

**Table 1:**
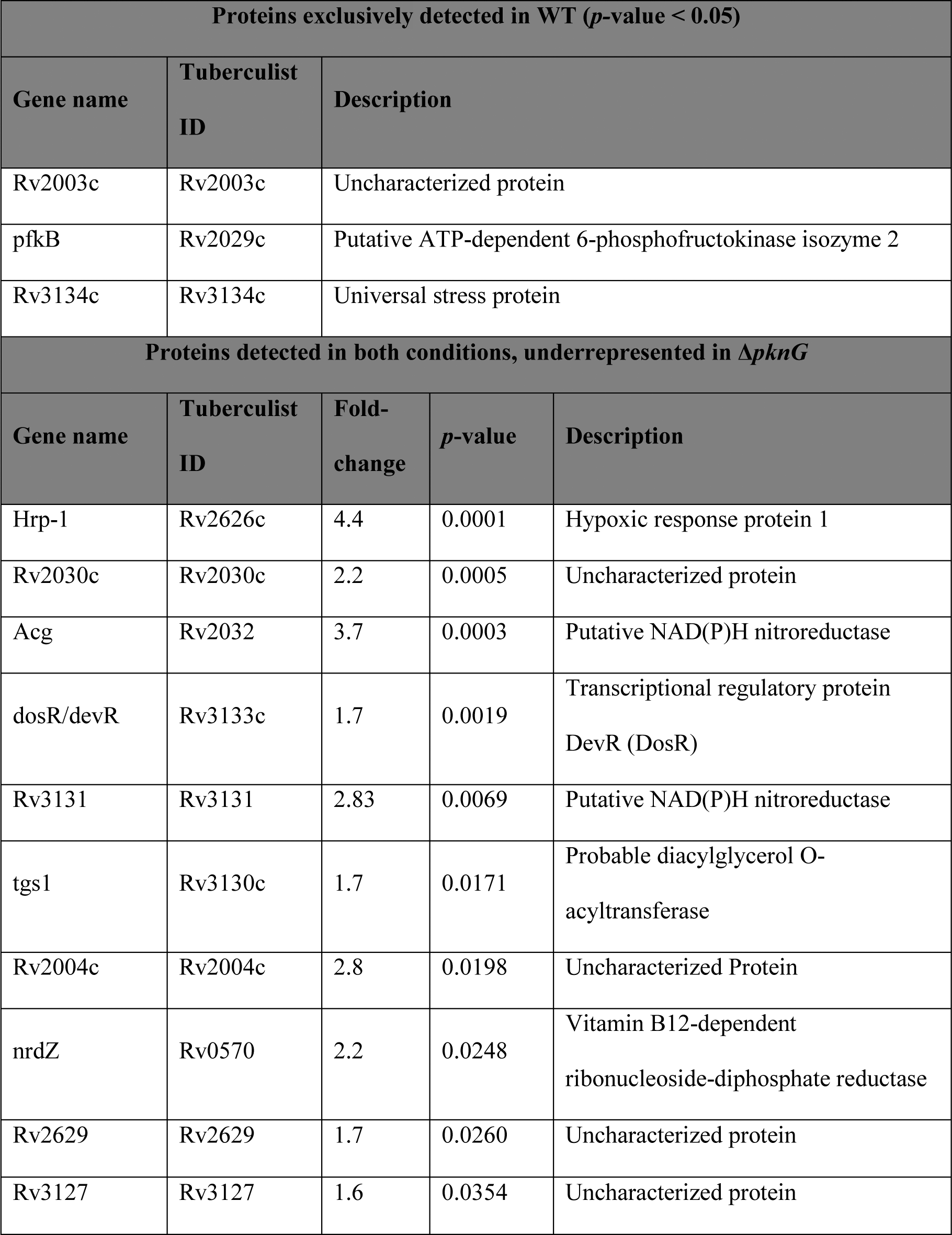
Proteins of DosR regulon statistically underrepresented in Δ*pknG*.

Remarkably, the proteins that showed the highest fold-changes among the differential protein dataset are enzymes involved in the synthesis of non-covalently attached outer layer lipids of the complex mycobacterial cell envelope (Table 2). The probable multifunctional mycocerosic acid synthase membrane-associated protein Mas was overrepresented in the WT strain (fold-change 19.9) while the mycolipanoate synthase Msl3 was overrepresented in the Δ*pknG* strain (fold-change 9.3). Besides, the protein N-acetylglucosamine-6-phosphate deacetylase NagA, involved in N-acetyl glucosamine utilization, is the protein that exhibited the highest fold-change among those overrepresented in the Δ*pknG* dataset (fold-change 22.5) (6). Also, many proteins that participate in mycolic acids biosynthesis were underrepresented in Δ*pknG* proteome, among them KasA, MmaA1, MmaA3, MmaA4, CmaA1, CmaA2, FbpB (Table 2 and Supplementary Table 5). In summary, our label-free LC-MS/MS analysis revealed that *pknG* deletion affects the relative abundance of enzymes involved in the biogenesis of the mycobacterial cell envelope, proteins involved in stress response mediated by TA systems and redox homeostasis. Moreover, PknG seems to directly or indirectly regulate the levels of expression of proteins that play essential roles in adapting *M. tuberculosis* to hypoxic conditions.

**Table 2:** Proteins involved in cell envelope metabolism showing statistically differential abundance between strains.

| Gene name | Tuberculist ID | Fold-change | *p*-value | Increased in: | Description |
| --- | --- | --- | --- | --- | --- |
| Mas | Rv2940 | 19.9 | 0.0002 | WT | Probable multifunctional mycocerosic acid synthase membrane-associated Mas |
| Msl3 | Rv1180/Rv1181 | 9.3 | 0.0006 | Δ*pknG* | Mycolipanoate synthase |
| NagA | Rv3332 | 22.5 | 0.0489 | Δ*pknG* | N-acetylglucosamine-6-phosphate deacetylase |
| mmaA3 | Rv0643c | 2.7 | 0.0066 | WT | Methoxy mycolic acid synthase |
| cmaA2 | Rv0503c | 2.1 | 0.0093 | WT | Cyclopropane mycolic acid synthase 2 |
| mmaA4 | Rv0642c | 1.6 | 0.0110 | WT | Hydroxymycolate synthase |
| FbpB | Rv1886c | 1.8 | 0.0132 | WT | Diacylglycerolacyltransferase/<br>mycolyltransferase Ag85B |
| cmaA1 | Rv3392c | 1.7 | 0.0176 | WT | Cyclopropane mycolic acid synthase 1 |
| mmaA1 | Rv0645c | 1.5 | 0.0233 | WT | Mycolic acid methyltransferase MmaA1 |
| kasA | Rv2245 | 1.6 | 0.0264 | WT | 3-oxoacyl-[acyl-carrier-protein]<br>synthase 1 |

Supporting this, functional enrichment analysis using differential proteins identified by 2D-DIGE and shotgun approaches indicated that the main biological process altered in *M. tuberculosis* Δ*pknG* is response to hypoxia (Supplementary Table 6). Also, consistent with the differential proteins discussed above, the biological processes “fatty acid biosynthetic process” and “oxidation-reduction” were statistically enriched in the WT proteome (Supplementary Table 6).

### Validation of differentially abundant proteins by parallel reaction monitoring (PRM)

To further confirm the relative abundance of some selected proteins we used a targeted proteomics strategy (PRM). After the refinement of peptide chromatograms using Skyline, we focused on two proteins that participate in the biosynthesis of lipids of the outermost layer of the cell envelope (Mas and Msl3) and 7 proteins involved in the response to hypoxia (Ald, Acg, DevR, Hrp-1, Icl, Lat, Rv2030c). We successfully validated the changes in the relative abundance of 5 of the 9 analyzed proteins (adjusted *p*-value < 0.05), namely Msl3, Mas, Hrp-1, Rv2030c, and Acg (Figure 4 and Figure 5). The quite large fold-changes recorded in the Δ*pknG* strain for Msl3 (2995 fold increment) and Mas (100 fold decrease) denoted a clear switch in enzymes responsible for the synthesis of free lipids in the outer layer of *M. tuberculosis* cell envelope. Among the proteins involved in response to hypoxia, Hrp-1 showed the highest expression reduction in the Δ*pknG* mutant (12.5 times) (Figure 4 and Figure 5). Thus, our results allow confirming that *pknG* deletion impacts on the outer lipids biosynthesis and the response to low-oxygen levels, two relevant physiological processes for the host-pathogen interaction and the bacterial survival within infected human cells.

**Figure 4:**
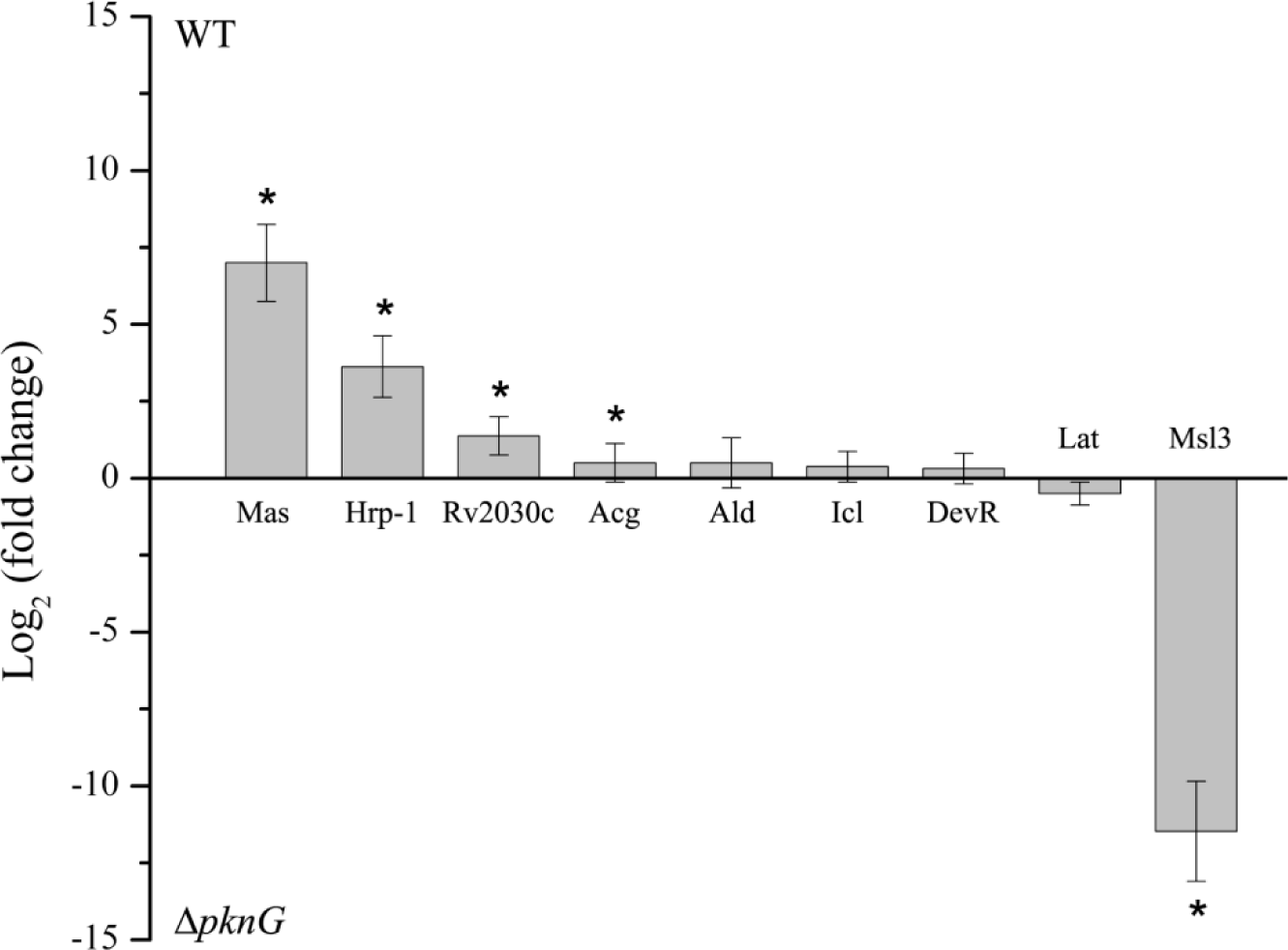
Validation of protein relative abundance changes by targeted proteomics (PRM). Bar graph showing the Log_2_(fold-change) of the proteins analyzed by PRM. The sum of transition areas was used as a quantitative measure and equalization to medians was employed for normalization. Positive values represent proteins increased in WT *M. tuberculosis* and negative values in the Δ*pknG* strain. Proteins showing statistically significant changes are indicated with an asterisk (adjusted *p-*value ˂ 0.05).

**Figure 5:**
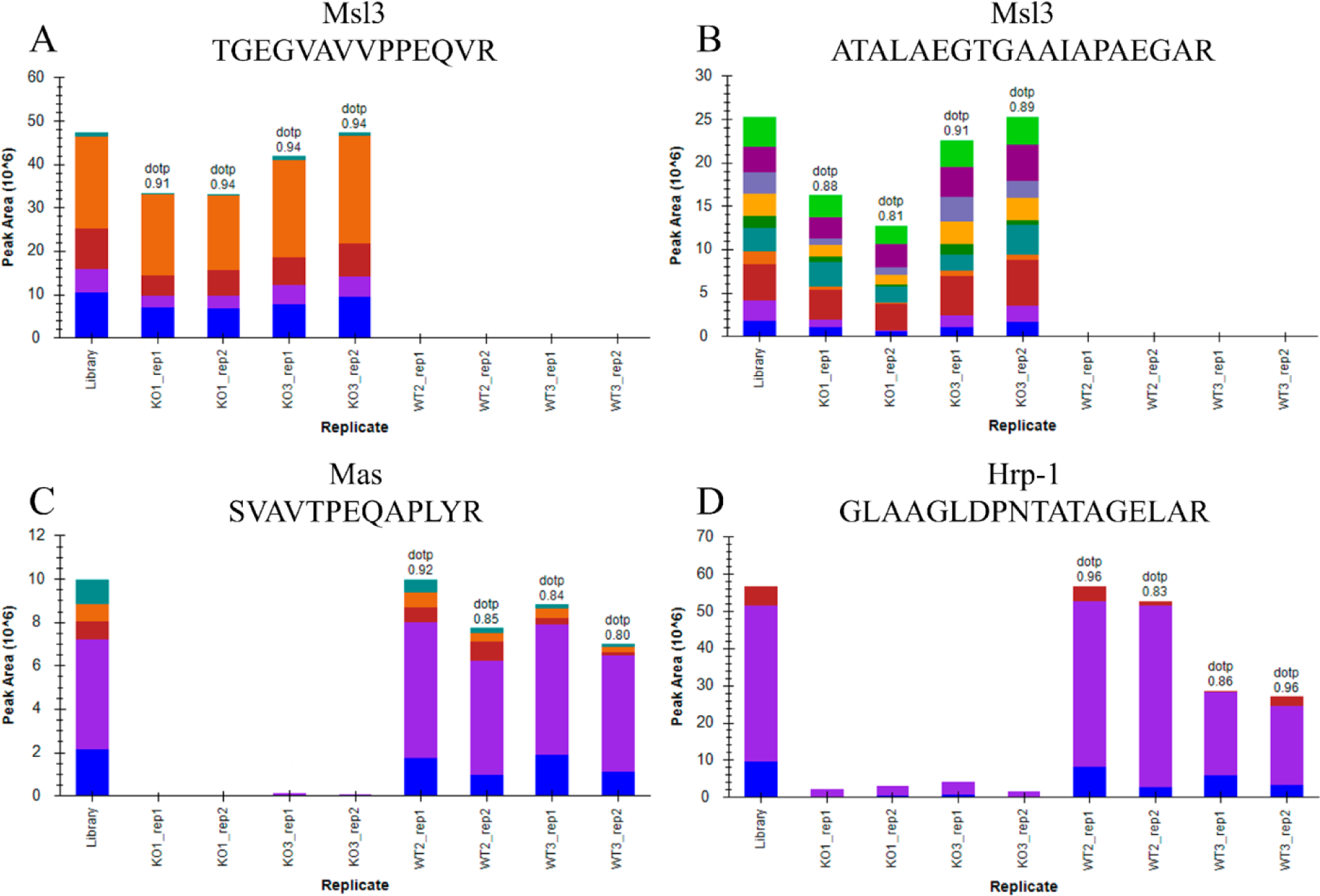
Changes in the relative abundance of selected peptides determined by PRM. Each bar graph shows the variation in abundance (represented as the sum of transition areas, each color in a bar represents an individual transition area). The dotp value is indicated. The sum of the transition area of each peptide in the library is also shown. Panel A and B: TGEGVAVVPPEQVR and ATALAEGTGAAIAPAEGAR peptides from Msl3 protein. Panel C: SVAVTEQAPLYR peptide from Mas protein. Panel D: GLAAGLDPNTATAGELAR peptide from Hrp-1 protein.

## DISCUSSION

In a previous work, we performed a 2D-DIGE analysis of WT and Δ*pknG M. tuberculosis* to identify substrates of PknG. Besides the differences in phosphorylation patterns, this analysis suggested that the strain lacking *pknG* had a more global proteomic change. In this work, we employed two comparative and quantitative proteomic approaches to assess the effect of *pknG* deletion on the total proteome of *M. tuberculosis*. Comprehensive proteomic profiling of *M. tuberculosis* Δ*pknG* indicated that as much as 15% of the detected proteins presented significant abundance changes compared to WT (5.7% of the predicted *M. tuberculosis* H37Rv proteins), pointing to a substantial remodeling of the proteome in the absence of PknG. Proteins with altered expression levels consistently mapped into defined biological processes relevant for virulence and bacterial survival within infected human cells.

During more than ten years, various research groups have contributed to an extensive phenotypic and functional characterization of mycobacteria lacking PknG, and the accumulated evidence clearly indicates that this kinase is crucial for pathogenicity. It was shown early that *M.tuberculosis* Δ*pknG* caused delayed mortality of highly susceptible infected mice, and presented decreased viability in an immunocompetent mice model (18). Furthermore, deletion of *pknG* has shown to impair granuloma formation in guinea pigs (20) and more recently a decreased capability of Δ*pknG* to resuscitate in a latent tuberculosis mouse model was reported (49). It was also demonstrated that the absence of PknG prevents mycobacterial survival within host macrophages (15) and leads to a growth defect under *in vitro* models of hypoxia (20). Finally, several pieces of evidence point to an altered cell envelope in the absence of PknG. Wolff *et al.* reported that a *Mycobacterium smegmatis* Δ*pknG* mutant strain presented severely altered cell surface properties, in terms of charge and hydrophobicity, and also showed that PknG is required for biofilm formation in several mycobacterial species, including *M. tuberculosis* and *Mycobacterium bovis* BCG (23). In addition, a diminished intrinsic resistance to antibiotics, possibly mediated by an altered permeability and hydrophobicity of bacterial cells, was reported for *M. smegmatis* Δ*pknG* (22).

Despite the overwhelming evidence supporting a role of PknG in growth inside the host and pathogenicity, the mechanism of action of this kinase at the molecular level is still a subject of extensive debate. While some studies showed that PknG regulation of glutamate and/or redox metabolism could play a key role for bacterial adaptation to the nutrient conditions found inside the host (19, 23, 50)(19), others proposed that the phosphorylation of host’s proteins is a key factor in PknG’s mediated virulence (15, 51). The proteomics changes reported here for Δ*pknG* are very consistent with the previously reported phenotypes, and shed some light on new molecular mechanism behind them.

Twenty-seven proteins were identified from 36 differential spots according to 2D-DIGE analysis; while the label-free LC-MS/MS approach allowed the detection of 228 additional differential proteins.

Seven differential proteins involved in response to hypoxia and lipid metabolism were detected using both approaches, with protein abundances changing in the same direction (Ald, BfrB, DesA2, FbpB, Icl, KasA, Rv0223c). Nineteen out of the 20 remaining differential proteins identified by DIGE were also detected in label-free LC-MS/MS experiments, but the differential levels could not be confirmed, either because proteins were identified with few spectra or because changes were not statistically significant. A possible explanation for this observation is that specific proteoforms, in particular phosphorylated ones, were responsible for the 2D-DIGE differential spots.

Very interestingly, the quite large proteomics changes associated with the deletion of this protein kinase mainly involves two relevant processes: cell envelope biogenesis and hypoxia response. These findings, discussed in detail below, allowed us to propose new mechanisms by which PknG mediates bacterial survival inside infected macrophages (Figure 6).

**Figure 6.**
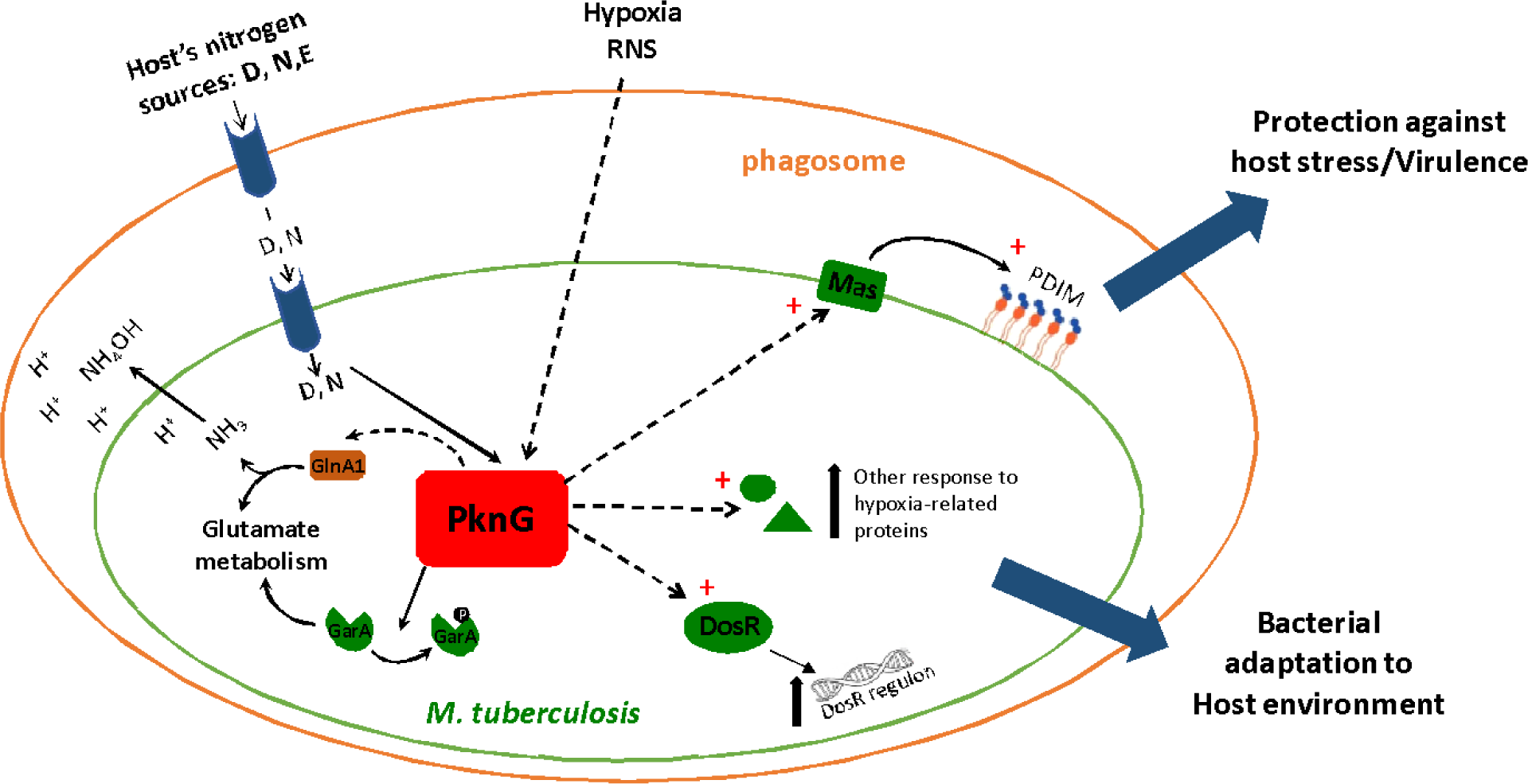
Model for the role of PknG in bacterial adaptation to nutritional stress inside the host. PknG responds to the nutrient availability in the host. Asp and Glu and Gln represent the principal source of nitrogen for intracellular bacteria (76) and were shown to activate PknG through the action of two accessory proteins, GlnH and GlnX (77); triggering the regulation of glutamate metabolism and nitrogen assimilation through GarA and possible GlnA1 phosphorylation (16, 17, 24). The results presented here allowed us to postulate that PknG also responds to low oxygen concentrations and directly or indirectly regulates the expression levels of DosR regulon as well as a group of proteins that are considered part of the proteomics hallmark of the hypoxic bacteria. In addition, PknG controls the levels of key proteins in the biosynthetic pathway of the virulence factor PDIM. The current model allows us to position PknG as a hub that redirects bacterial metabolism in response to the availability of nutrients, including oxygen, thus facilitating bacterial survival within the host.

### Cell envelope biosynthesis

Our results indicated that the expression levels of proteins implicated in the biosynthesis of different components of the *M. tuberculosis* cell envelope were altered in Δ*pknG.* Proteins involved in peptidoglycan component recycling and cell wall synthesis were overrepresented in Δ*pknG*. NagA, an enzyme that catalyzes a critical step for the synthesis of peptidoglycan precursors and its recycling (52) was 22.5 fold enriched in Δ*pknG* (Table 2). In addition, several proteins located in an operon related to peptidoglycan synthesis, cell growth and shape (the STPKs PknA and PknB, and the protein FhaA) (53, 54), were also overrepresented in the Δ*pknG* dataset (Supplementary Table 5). On the contrary, the biosynthetic pathway of other distinctive components of the cell envelope of these bacteria, mycolic acids, is underrepresented in Δ*pknG* (Table 2 and Supplementary Table 5). Altogether, these observations suggest that PknG could be involved in the regulation of the structure of cell wall core of *M. tuberculosis*.

However, the most important proteomic changes are related to the biosynthetic pathways of other lipid components of the cell envelope. Our results strongly suggest a major change in the composition of the bioactive complex lipids found in the outermost layer of the cell envelope. Two enzymes of the polyketide synthase family that participates in the biosynthesis of branched fatty acids, the Multifunctional mycocerosic acid synthase membrane-associated Mas and the Mycolipanoate synthase Msl3, presented high fold-changes in Δ*pknG*. These impressive fold-changes in opposite directions were further confirmed by PRM targeted proteomics, showing a 100 fold decrease of Mas levels and 2995 fold increase in Msl3 levels in Δ*pknG*. Mas and Msl3 share the same enzymatic activity and have substantial sequence identity, but participate in different biosynthetic pathways (55). Mas is involved in the biosynthesis of the cell envelope’s phthiocerol dimycocerosates (PDIMs), which are unique for slow growing mycobacteria and have a key role in *M. tuberculosis* pathogenesis (5, 55–57). On the other hand, Msl3 is a Mas-like enzyme involved in the biosynthesis of critical constituents of polyacyltrehaloses (PATs), another kind of free lipids of the cell envelope of *M. tuberculosis* that are also found exclusively in virulent strains (55, 58). However, in contrast to PDIMs, there is experimental evidence showing that PATs are not essential for virulence (55, 58). The Mas enzyme is required for full virulence of *M. tuberculosis* and it was shown to participate in lipid synthesis during infection (59, 60). The products of the biosynthetic pathway in which Mas participates, PDIMs, are one of the very early reported virulence factors of *M. tuberculosis* (61) and have an important role in the fate of the bacteria inside infected cells. Indeed, disruption of genes involved in PDIMs biosynthesis led to strains unable to inhibit phagosome acidification and maturation (62), a phenotype already described for Δ*pknG* (15). Although the mechanisms underlying PDIMs–mediated virulence are still not fully understood, a role in the modulation of the immune response, the properties of cell surface and the protection against nitrogen reactive species has been reported for this cell envelope component (63, 64).

Altogether, these observations allow us to postulate that there is a switch in the type of free lipids synthesized by Δ*pknG* in *M. tuberculosis*, with increased levels of PATs and decreased levels of the PDIMs virulence factors. In this scenario, it is tempting to speculate that the much lower levels of Mas, and possibly of its biosynthetic products PDIMs, could contribute to the observed altered physicochemical properties of the cell envelope and the growth defects inside host macrophages as previously reported for Δ*pknG*.

### Response to hypoxia

The evidence arising from both, label free LC-MS/MS and DIGE experiments, consistently indicated that proteins involved in the response to hypoxia were downregulated in Δ*pknG*. On one hand, around one third of the underrepresented protein spots in DIGE gels of Δ*pknG* are increased in hypoxia models of *M. tuberculosis* (37, 65) (Supplementary Table 3). Shotgun analysis further supported these results revealing that response to hypoxia is the main biological process altered in Δ*pknG*. Adaptation to low oxygen conditions in mycobacteria is mainly mediated by the DosR regulon, which comprises around 50 genes that are up-regulated by the DosR response regulator under oxygen limitation conditions (12, 66, 67). Our proteomics results allowed us to detect several proteins from this regulon as underrepresented in Δ*pknG* (Table 1). In addition to DosR regulon proteins, several proteins known to be relevant for the entry in the hypoxic non-replicative state were also underrepresented in Δ*pknG*: Ald, Icl, BfrB, Tuf, Lat. All of these differential proteins are thought to be involved in the adaptation of the bacteria to hypoxic conditions and other stress conditions found inside macrophages and granulomas, and are jointly considered a distinctive proteomic hallmark of the mycobacterial response to hypoxia (38–41, 66). Only one of the proteins of this hypoxic proteomics fingerprint could not be detected with altered levels in Δ*pknG*: the HspX protein. HspX and Hrp-1constitute the most paradigmatic DosR regulon induction markers and in turn are the proteins whose levels change most dramatically in response to hypoxia (68, 69). Interestingly enough, while Hrp-1 levels were very significantly decreased in Δ*pknG*, HspX is among the proteins whose global levels were not altered in shotgun analysis, but presented differential spots in 2D-DIGE. This intriguing finding deserves further investigation. One possibility is that PknG, through phosphorylation of specific substrates, may add an additional level of control on the DosR regulon, allowing to differentially tune the levels of its various components. In fact, the regulation of DosR activity by Ser/Thr protein kinases (in addition to the well-studied phosphorylation by His protein kinases) has already been shown. On one hand, phosphorylation of Thr198 and Thr205 by PknH contributes to the DosR dimerization and enhances its transcriptional activity (73). On the other hand, phosphorylation of DosR Thr180 by overexpression of the catalytic domain of PknB in *M. smegmatis* negatively affected the DNA-binding affinity of the regulator to its target DNA sequence (70). Thus, an attractive hypothesis is that a defect in DosR phosphorylation in Δ*pknG* could be mediating the low levels of hypoxic response proteins. In fact, published data support this hypothesis. Bae *et al.* evaluated the interaction between DosR and the 11 STPKs codified by *M. tuberculosis* using yeast two-hybrid assay, and PknG showed the strongest interaction (70). However, these authors use HspX as a reporter for DosR induction, and this led them to dismiss the possible physiological relevance of the PknG-DosR interaction. Based on our results, HspX is not a good marker of a PknG-mediated induction of hypoxia response. The possible direct phosphorylation of DosR by PknG deserves to be investigated.

Altogether, 19 out of 59 proteins exclusively detected in WT, and 38 out of 92 proteins overrepresented in this same strain, have previously been reported to be up-regulated in the proteome of hypoxic bacteria (4, 14, 38, 41, 66, 71–73). This is in very good agreement with reports showing that mycobacteria lacking PknG were unable to grow under hypoxic conditions (50). These authors showed that this effect was mediated by GarA phosphorylation. The proteomic analyses reported here points to the PinG’s direct or indirect control of the expression levels of key proteins for hypoxic lifestyle switch as an additional mechanism behind Δ*pknG*’s inability to grow under hypoxia. It is interesting to note that the conserved kinase domain of PknG is flanked by an N-terminal rubredoxin-like domain composed by an iron ion coordinated to four conserved cysteine residues (74, 75). These domains typically participate in electron transfer reactions, and in the case of PknG a role in catalysis regulation has been demonstrated for the rubredoxin-like domain (75). The possible participation of this rubredoxin like domain in the direct sensing of low oxygen concentrations is an interesting hypothesis to be investigated.

Altogether, our proteomic data showed that the deletion of *pknG* gene from *M. tuberculosis* causes an alteration in the relative abundance of many proteins that participate in cell envelope biosynthesis, adaptation to hypoxic conditions and the establishment of a persistent bacterial state, in fully agreement with previous functional knowledge about Δ*pknG*. These results, together with previously published data that indicated that PknG sense amino acid availability and regulates glutamate metabolism, allowed us to start delineating a model for PknG’s regulation of bacterial adaptation to the nutritional conditions found in the host (Figure 6).

Early macrophage infection studies indicated that deletion of PknG in pathogenic mycobacteria leads to its rapid degradation in mature lysosomes, and suggested that host protein phosphorylation upon PknG secretion could be the mechanism of action (15). Instead, the results presented here support the idea that the direct or indirect regulation of the expression levels of a group of proteins that are relevant for the adaptation of the bacterium to the host environment and the induction of a mycobacterial persistent state might account in part for the effect of PknG on bacterial survival inside the host.

## Supporting information

Protein identification LC-MS/MS-Additional information

Differential proteins-LC-MS/MS

Biological process enrichment

Total identified proteins-LC-MS/MS

PRM isolation list

Protein identification-DIGE

## Abbreviations

DIGE: Difference In-Gel Electrophoresis
DosR: Dormancy Survival Regulator
FDR: false discovery rate
Hrp-1: Hypoxic Response Protein 1
Mas: Probable multifunctional mycocerosic acid synthase membrane-associated
Msl3: Mycolipanoate synthase
PAT: polyacyltrehaloses
PDIM: phthiocerol dimycocerosate
PRM: parallel reaction monitoring
STD: internal standard
STPK: Ser/Thr protein kinase
TA: toxin-antitoxin
TB: tuberculosis
TCS: two-component system

## ACKNOWLEDGMENTS

This work was supported by grants from Agencia Nacional de Investigacion e Innovacion, Uruguay (ANII, FCE_3_2013_1_100358 and FCE_1_2014_1_104045) and FOCEM - Fondo para la Convergencia Estructural del Mercosur (COF 03/11). MG and BR were supported by fellowships from ANII [POS_NAC_2012_1_8824, POS_NAC_2015_1_109755, POS_FCE_2015_1_1005186]. AC and RB were supported by the Agence Nationale pour la Recherche (France). The authors would like to thank Dr. Av-Gay for kindly providing us with the *M. tuberculosis* Δ*pknG* strain.

## Figure Legends

**Supplementary Figure 1:**
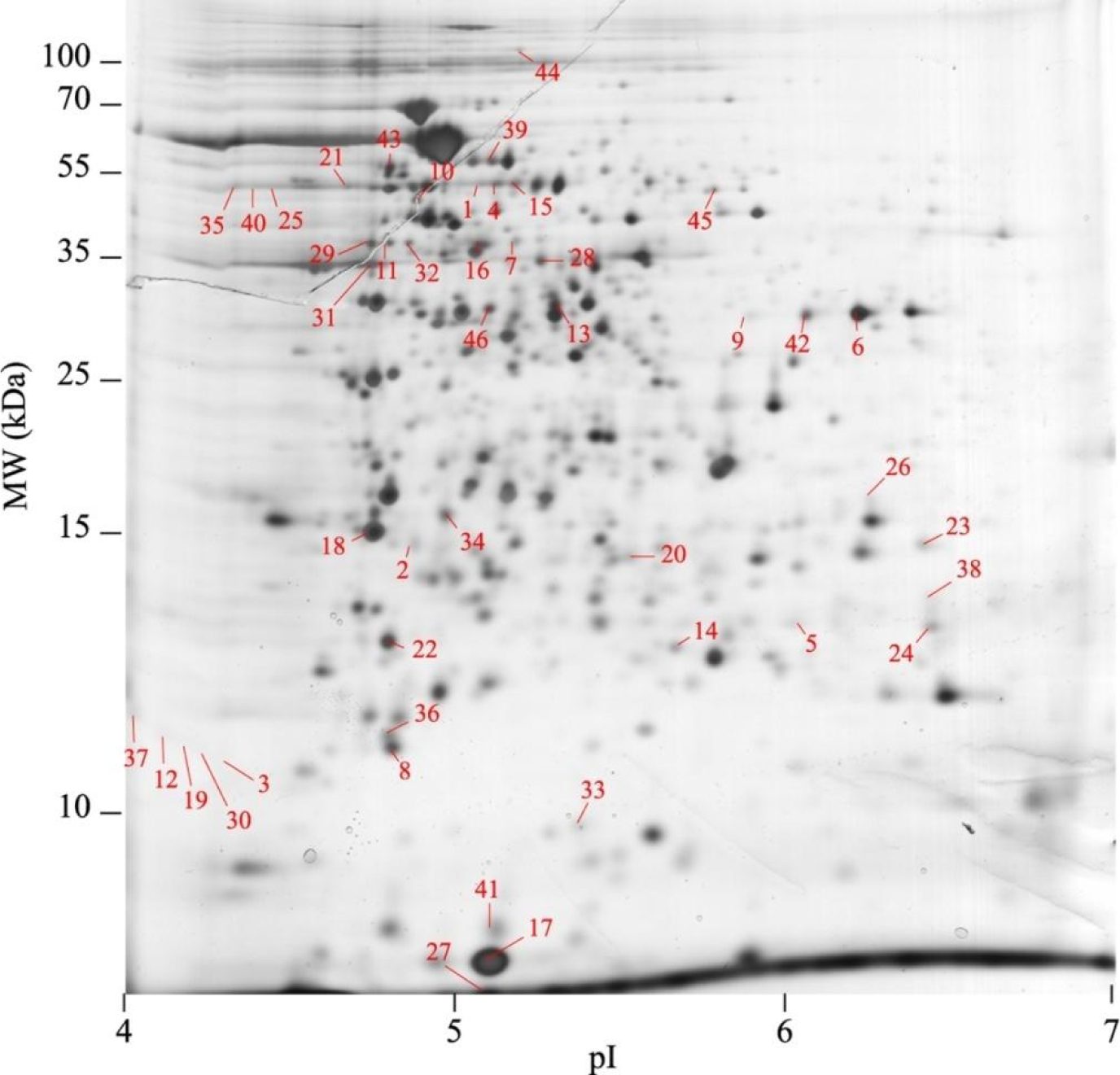
Spot numbers of protein spots selected after 2D-DIGE analysis and identified by MALDI-TOF/TOF MS. GarA and GlnA1 spots were previously identified (spots numbers 1, 3, 4, 12, 15, 19, 21, 25, 30, 35, 37, 40) (24).

**Supplementary Figure 2:**
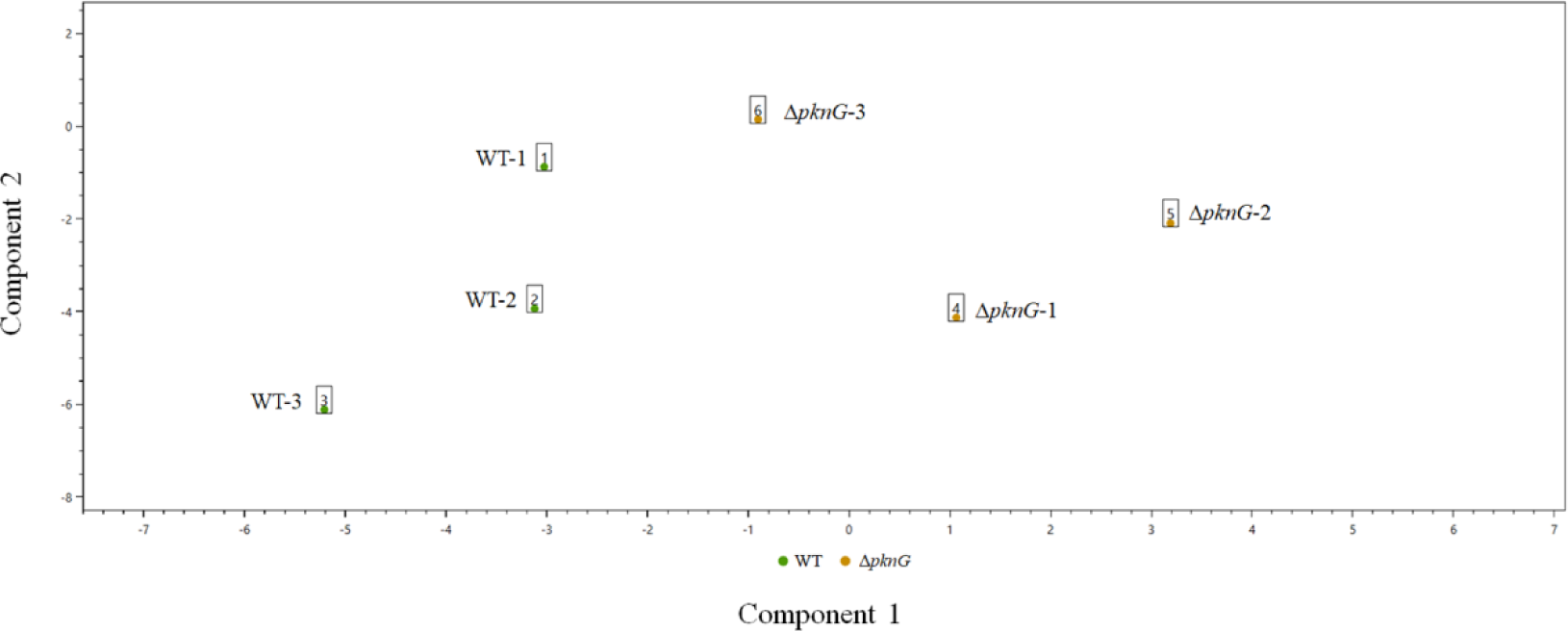
Principal Component Analysis of WT and Δ*pknG* sample sets. Each green spot represents a WT proteomic dataset. Each orange spot represents a Δ*pknG* dataset.

**Supplementary Figure 3:**
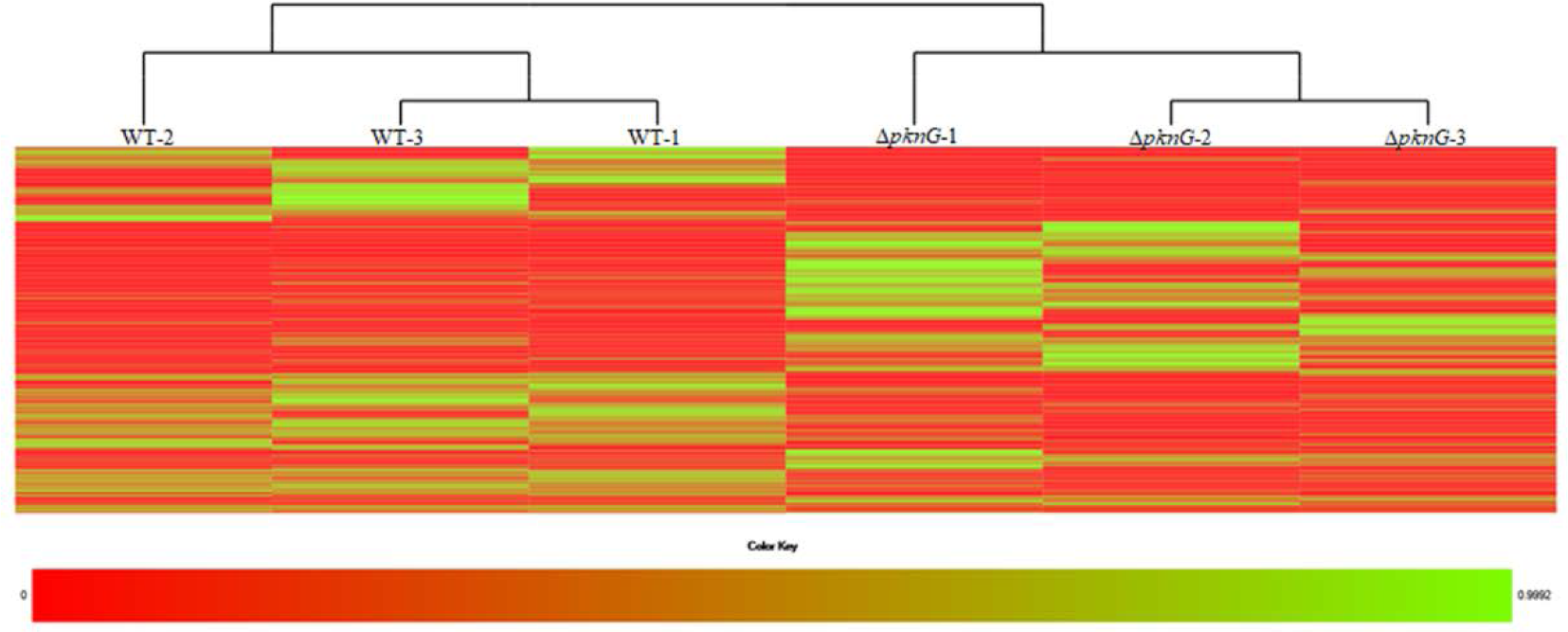
Clustergram analysis visualized as a heat map. Each column represents a dataset. Each row shows an individual protein. The gradient color indicates the relative abundance from 0 (red) to 1 (green).

**Supplementary Table 1**: Total of proteins identified by LC-MS/MS analysis. Each tab corresponds to each strain replicate.

**Supplementary Table 2**: PRM isolation list peptides.

**Supplementary Table 3**: Protein spots selected after 2D-DIGE analysis and identified by MALDI-TOF/TOF MS. Spots numbering corresponds to Supplementary Figure 1.

**Supplementary Table 4**: Complementary data of LC-MS/M identification.

**Supplementary Table 5**: Differential proteins by label free LC-MS/MS analysis. Proteins uniquely detected in one of the two conditions were determined using the PatternLab’s Venn Diagram module (*p* - value < 0.05). Differentially abundant proteins were determined using the PatternLab’s T-Fold module.

**Supplementary Table 6**: Details of the Biological Process Enrichment test performed with Panther Sever.

